# Hidden causal inference delineates dynamic lncRNA regulation in autism spectrum disorder

**DOI:** 10.64898/2026.01.25.701433

**Authors:** Junpeng Zhang, Xuemei Wei, Chunwen Zhao, Zidan Hu, Shujie Fan

**Affiliations:** School of Engineering, Dali University, Dali, Yunnan, 671003, China

**Keywords:** lncRNA, Single-nucleus RNA sequencing, Causal inference, Dynamic lncRNA regulation, Autism spectrum disorder

## Abstract

Long non-coding RNAs (lncRNAs) are increasingly implicated in autism spectrum disorder (ASD), yet their causal regulatory mechanisms remain poorly characterized. Conventional static causal inference methods fail to capture the dynamic interplay of lncRNAs across diverse brain biological contexts. Here, we develop a hidden causality-based method Clean to infer dynamic lncRNA causal regulation in ASD. Our analysis reveals that dark causality, representing a hybrid form of positive and negative interdependencies, dominates lncRNA-mediated regulation in ASD. Moreover, dynamic lncRNA causal regulation emerges in diverse brain biological contexts, and 14 ASD risk lncRNAs significantly contribute to disease pathogenesis through disrupted gene regulatory programs. We also illustrate that lncRNA expression signatures enable robust classification of sex and cell type, offering potential diagnostic lncRNA biomarkers. Notably, 20 immune-related lncRNAs are prominently involved in the immune, suggesting a potential role for neuroimmune regulation. Additionally, cell-cell interaction networks diverge substantially between ASD and Normal cohorts. This study establishes a paradigm for delineating the dynamic causal regulation of lncRNAs, underscoring their diagnostic and therapeutic potential in ASD.

## Introduction

Autism spectral disorder (ASD) is a group of neurodevelopmental conditions characterized by two core deficits, including impairments in verbal and nonverbal communication, and restricted/repetitive patterns of behaviour, interests and activities (Rosen et al. 2021). Although the etiology of ASD remains unclear, current consensus implicates that it is caused by a combination of genetic and environmental factors. These genetic and environmental triggers induce cellular dysfunction, leading to neuroinflammation (Madore et al. 2016), excitation/inhibition imbalance (Hollestein et al. 2023), and white matter abnormalities (Ohta et al. 2020). Traditional ASD diagnosis relies on clinical symptoms, including social communication, social interactions, and behavioral/sensory processing. However, these clinical symptoms demonstrate limited applications across sex, age, and intellectual ability. Furthermore, core clinical symptoms (e.g., impaired social communication, restricted behaviours) are emerged clearly only after age three, delaying early intervention opportunities. Emerging diagnostic methods leverage ASD susceptibility genes as biomarkers for ASD risk prediction. These susceptibility genes frequently harbor mutations driven by sex-linked modifiers, epigenetic regulation, copy number variations, environmental exposures or double-hit mutational events (Iakoucheva et al. 2019a). Biomarker-based molecular diagnosis is unaffected by age, sex, or intellectual ability, facilitating earlier detection and intervention (Iakoucheva et al. 2019a; Frye et al. 2019; Zhuang et al. 2024). This paradigm shift toward molecular diagnosis holds promise for improving ASD intervention outcomes.

Generally, ASD biomarkers are predominantly limited to protein-coding genes, as exemplified by the SFARI (Abrahams et al. 2013), AutismKB (Yang et al. 2018), and AutDB (Pereanu et al. 2018) databases. Recent studies, however, reveal critical roles for non-protein-coding genes in ASD pathogenesis (Ghafouri-Fard et al. 2022). For example, the RNADisease database (Chen et al. 2023) systematically documents non-coding RNAs (ncRNAs) related to ASD. These ncRNAs modulate gene expression through DNA-level, pre-transcriptional, transcriptional, post-transcriptional, translational, and post-translational processes, thereby contributing to ASD progression (Dominguez-Alonso et al. 2023). Among the diverse classes of ncRNAs, long non-coding RNAs (lncRNAs), defined as RNA transcripts exceeding 200 nucleotides, have emerged as important regulators in ASD (Ghafouri-Fard et al. 2022). Emerging evidence suggests that several lncRNAs (e.g., *H19, NEAT1, MALAT1, MEG3, PVT1, MIR600HG, CSNK1A1P1, PCAT-1, PCAT-29, lincRNA-ROR, LINC-PINT, lincRNA-p21*) serve as promising non-invasive biomarkers by modulating synaptic plasticity and neurodevelopmental pathways (Bernard et al. 2010; Taheri et al. 2021; Jiang and Chen 2023; Jiang et al. 2023; Rahmani et al. 2024; Sane et al. 2024; He et al. 2025; Li et al. 2025). Investigations into lncRNA-mediated regulation offer novel insights into the molecular pathogenesis underlying ASD, and contribute to discovering novel lncRNAs as therapeutic targets for ASD (Srinivas et al. 2023). Nevertheless, the functional characterization of lncRNAs in gene regulation and their clinical potential as ASD biomarkers remains underexplored.

Since lncRNAs are characterized by dynamic regulation, low expression, and low conservation, they exhibit a great diversity in gene regulatory mechanisms (Chen and Kim 2024; Ferrer and Dimitrova 2024). In ASD, the mechanistic complexity of gene regulation suggests that the lncRNA regulation is inclined to be dynamic across various conditions. However, current approaches generally fail to capture conditional shifts in lncRNA regulation, limiting insights into how condition-specific perturbations drive ASD pathogenesis. While Cycle (Xiong et al. 2024) enables dynamic modelling of lncRNA regulatory networks across diverse cell types in ASD, it remains restricted to inferring cell-type-resolved correlative regulation, neglecting other ASD biological contexts (e.g. diagnosis, region, age and sex) and causal regulation by lncRNA. However, such as changes in diagnosis, region, age and sex, most ASD mechanisms can be directly affected, and the dependency of lncRNA causal mechanisms on such biological contexts should be learned. To address this gap, we develop a hidden causality-based method named Clean (dynami<u>C</u> causal r<u>e</u>gul<u>a</u>tio<u>n</u>), to delineate dynamic causal regulation by lncRNAs across diverse ASD biological contexts, such as diagnosis, region, age, sex, and cell type. For a comprehensive understanding of lncRNA causal regulation in ASD, Clean infers three types of causal relationships mediated by lncRNAs: positive, negative, and dark causality (a hybrid form of positive and negative interdependencies). Our analysis reveals that a dominant “dark causality” governs lncRNA-mediated regulation, identifies 14 ASD risk lncRNAs that disrupt gene regulatory programs, and demonstrates that potential lncRNA biomarkers can robustly classify sex and cell type. Furthermore, we have identified 20 immune-related lncRNAs, implicating a role for neuroimmune regulation, and observed a substantial alteration in cell-cell interaction networks between ASD and Normal cohorts. Collectively, Clean would provide a foundation for stratifying ASD etiologies, and offering technical support for precision therapeutics tailored to dynamic causal regulation states.

## Results

### Dynami<u>C</u> causal r<u>e</u>gul<u>a</u>tio<u>n</u> (Clean)

We introduce Clean, a hidden causality approach to elucidate dynamic lncRNA causal regulation in ASD from snRNA-seq data (**Fig. 1**). According to the diagnosis (ASD and Normal, 2 data slices), region (ACC and PFC, 4 data slices), age (<18 and >=18, 4 data slices), sex (Male and Female, 4 data slices) and cell type (17 cell types, 34 data slices) of brain cells, Clean divides the snRNA-seq data into 48 data slices (**Supplemental Data 1**). Since most of highly similar brain cells correspond to technical noise and not to biologically relevant heterogeneity, Clean builds metacells to remove technical noise while preserving the biological information of each data slice. Here, metacells are disjoint homogeneous groups of highly similar cells (Bilous et al. 2024). To investigate state changes in physiological and pathological conditions, Clean further uses trajectory inference to order brain metacells of each data slice along a linear trajectory. To explore dynamic causal regulation by lncRNAs, Clean employs hidden causal inference to identify causal relationships between lncRNAs and mRNAs in each data slice. Finally, Clean performs heterogeneity analysis to delineate dynamic lncRNA causal regulation across diverse brain biological contexts (i.e. diagnosis, region, age, sex and cell type).

**Figure 1.**
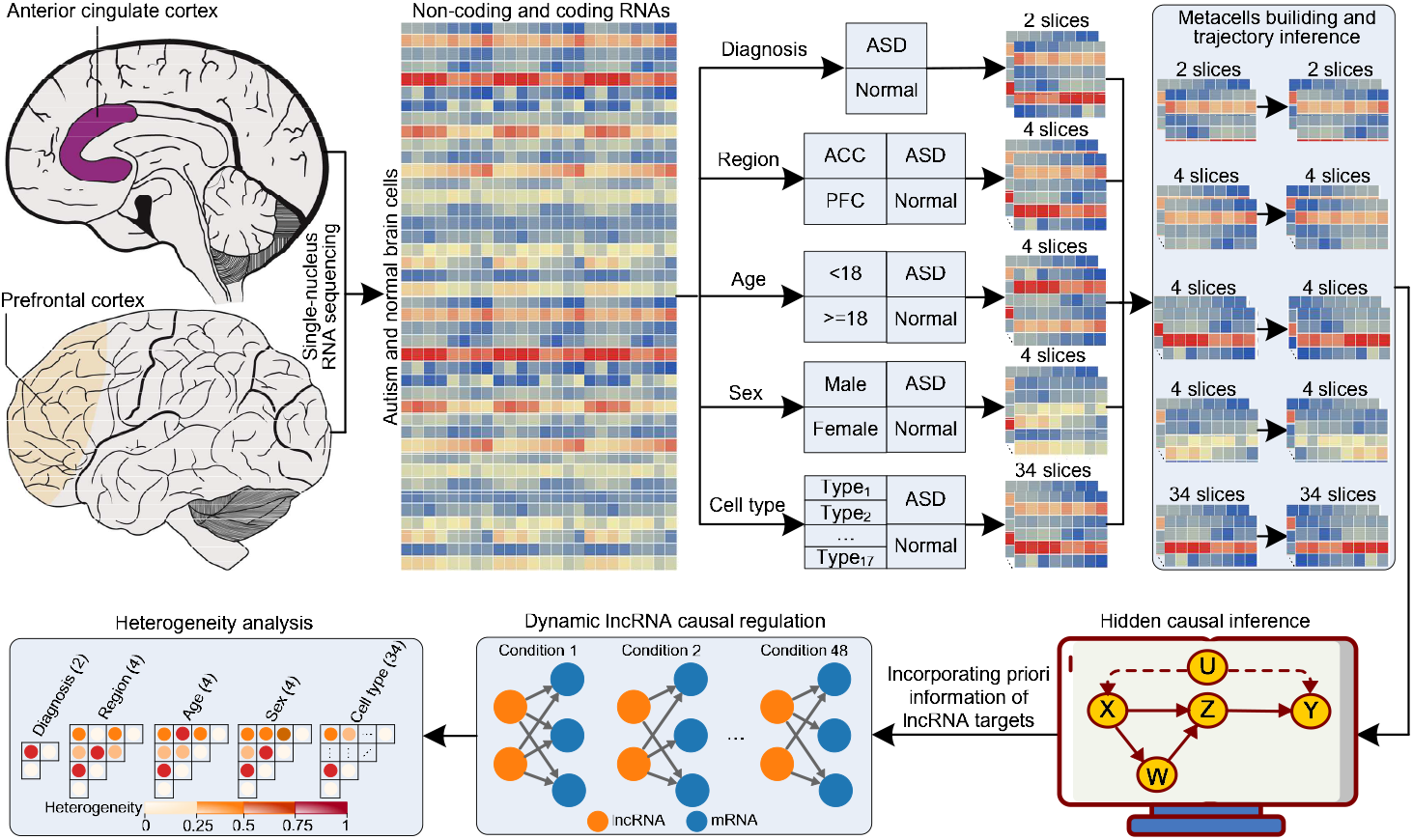
Schematic workflow of Clean. According to brain biological contexts (i.e. diagnosis, region, age, sex and cell type), the snRNA-seq data of autism and normal brain cells is splitted into 48 data slices. For each data slice, metacells are built for removing technical noise and preserving the biological information of it. For the identified metacells, Clean uses trajectory inference to order them along a linear trajectory. Given the processed 48 data slices, Clean applies hidden causal inference and incorporates priori information of lncRNA targets to infer dynamic lncRNA causal regulation. Based on the identified dynamic lncRNA causal regulation, Clean further conducts heterogeneity analysis to capture conditional shifts in lncRNA regulation.

### Dark causality emerges as the dominant type in lncRNA regulation across brain biological contexts

Causality is an intrinsic, structural property of biological networks, governing the fundamental landscape of a biological system (Barabási and Oltvai 2004). To characterize the spectrum of lncRNA causal regulation across diverse brain biological contexts including diagnosis, region, age, sex, and cell type, we quantify the prevalence of three distinct causal types: positive, negative, and dark causality. We find that dark causality, a hybrid form of positive and negative interdependencies, is the dominant causal type throughout all biological contexts examined, and this pattern remains robust regardless of whether priori information is incorporated into the causal inference (**Fig. 2, Supplemental Data 2**). The invariance of dark causality across diagnosis, region, age, sex, and cell type indicates that it is a fundamental and pervasive mode of lncRNA regulation in ASD, suggesting that lncRNA-mediated gene regulation often operates through complex, blended influences rather than purely positive activation or negative repression.

**Figure 2.**
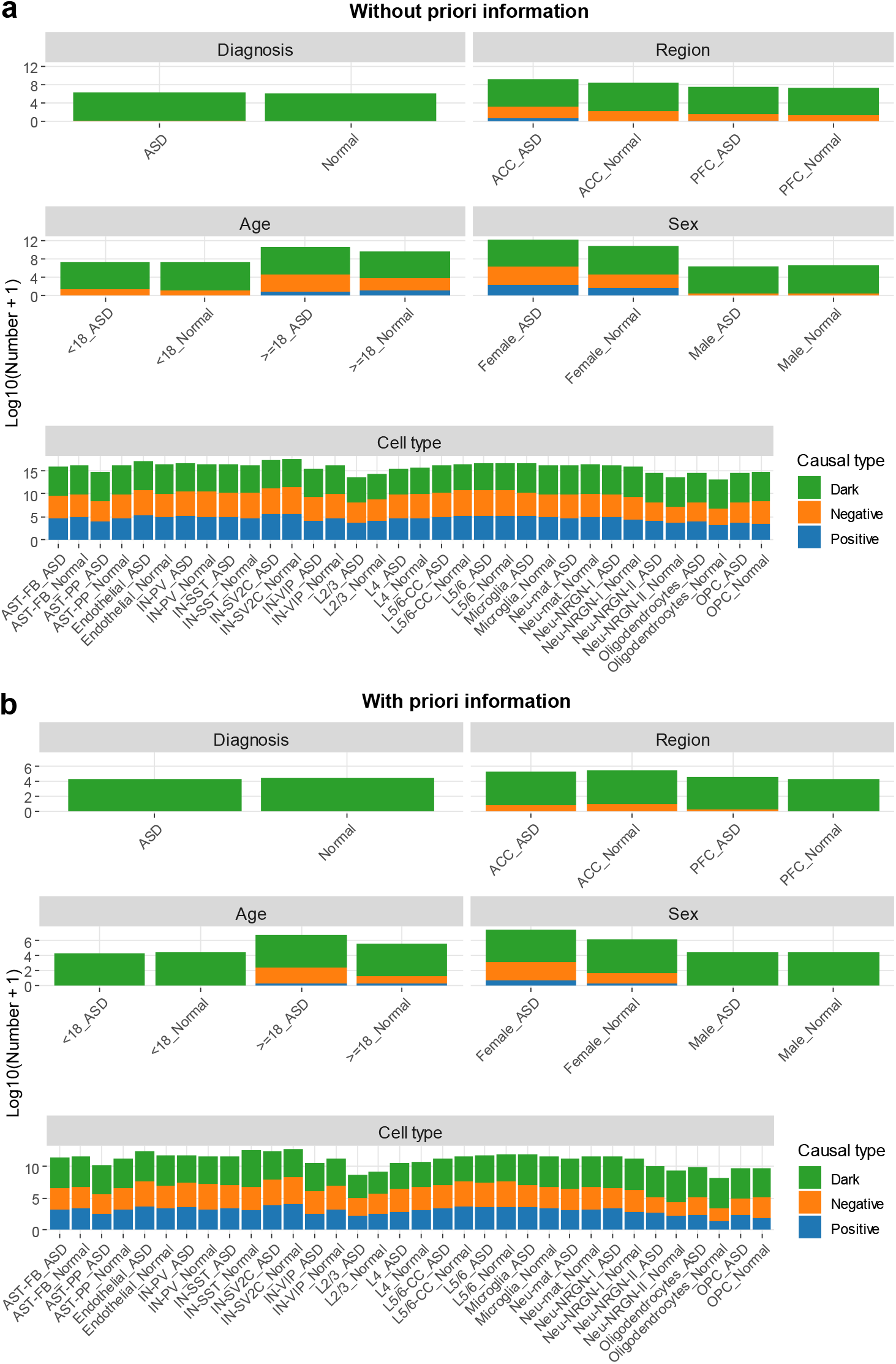
The spectrum of lncRNA causal regulation across diverse brain biological contexts. Number of positive, negative and dark causality (a) without and (b) with priori information in diverse brain biological contexts including diagnosis, region, age, sex, and cell type.

### Dynamic lncRNA causal regulation emerges in diverse brain biological contexts

It is demonstrated that lncRNAs exhibt dynamic and context-dependent regulatory roles in ASD, contributing to the disorder’s etiological complexity (Velmeshev et al. 2019; Parikshak et al. 2016). To investigate the dynamic nature of lncRNA-mediated regulation, we analyze the scale and heterogeneity of lncRNA causal regulatory networks across diverse brain biological contexts, including diagnosis, region, age, sex, and cell type (**Fig. 3**). For diagnosis, the network is substantially expanded in ASD compared to Normal, with a heterogeneity score of 0.40, indicating both quantitative increase and structure rewiring of regulatory interactions (**Fig. 3a**). Between brain regions, the number of regulatory relationships differs markedly, accompanied by heterogeneity scores >0.30 between ACC and PFC, reflecting strong spatial compartmentalization of network connectivity (**Fig. 3b**). In terms of age, we observe a clear shift in the quantity of regulatory interactions between donors under and over 18 years old, alongside heterogeneity scores >0.31, suggesting developmental stage-dependent network reorganization (**Fig. 3c**). For sex, sex-dimorphic differences are evident in both the number of causal interactions and heterogeneity scores, supporting sex as a key layer of variation in regulatory architecture (**Fig. 3d**). At the cell-type level, profiling reveal striking differences in both the number and heterogeneity of causal relationships, with immune cells such as Microglia in ASD showing particularly high connectivity and distinct regulatory patterns (**Fig. 3e**). Together, these results establish that lncRNA causal regulatory networks are dynamically organized in a multi-layered, context-dependent manner, varying in both scale and structural heterogeneity across biological contexts.

**Figure 3.**
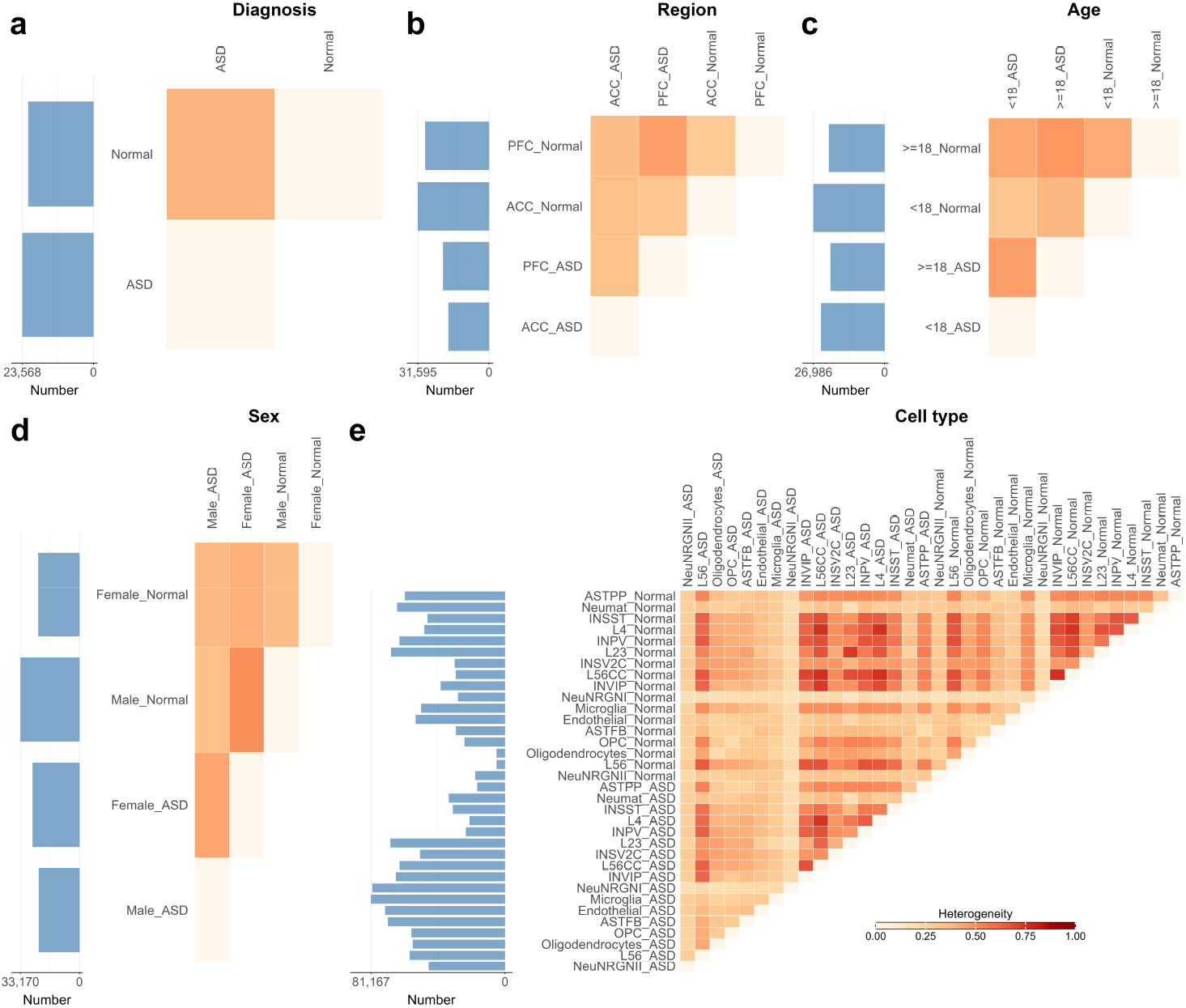
The scale and heterogeneity of lncRNA causal regulation. The lncRNA causal regulation across diverse brain biological contexts, including (a) diagnosis, (b) region, (c) age, (d) sex, and (e) cell type. The bar plots show the number of lncRNA-mRNA causal relationships in each condition, and the heat maps indicate the heterogeneity score of lncRNA causal networks between conditions in each biological context.

### ASD risk lncRNAs are significantly involved in the pathogenesis of ASD

In ASD, large-scale genetic and transcriptomic studies have revealed widespread dysregulation of lncRNA expression and highlighted their genomic loci as enriched for ASD-associated risk variants (Parikshak et al. 2016; Satterstrom et al. 2020). To systematically investigate the potential role of lncRNAs in the molecular etiology of ASD, we perform differential causal network analysis between ASD and Normal, which lead to the identification of 14 ASD risk lncRNAs (*TRMT2B-AS1, MFSD12-AS1, LINC00903, EPC1-AS1, LINC01632, LINC01840, P3H2-AS1, HRG-AS1, TCF4-AS1, MYCBP2-AS1, WDFY3-AS2, RNPC3-DT, MELTF-AS1, ZNF236-DT*) exhibiting statistically significant risk scores (**Supplemental Data 3**). As illustrated in **Fig. 4a**, the 14 ASD risk lncRNAs exhibit broad enrichment across multiple functional categories, including BP, MF, CC, KEGG, and Reactome terms (**Supplemental Data 4**). Among the 14 ASD risk lncRNAs examined, *TRMT2B-AS1, MFSD12-AS1, EPC1-AS1* and *LINC00903* show the highest degree of enrichment for functional annotation terms. Moreover, all 14 identified ASD risk lncRNAs are collectively enriched in at least one functional term directly linked to neurodevelopment and synaptic function, which are known to underlie ASD pathogenesis (**Fig. 4b**). Collectively, these results imply that the identified 14 ASD risk lncRNAs play a significant functional role in the molecular pathogenesis of ASD, and may offer promising targets for further mechanistic investigation.

**Figure 4.**
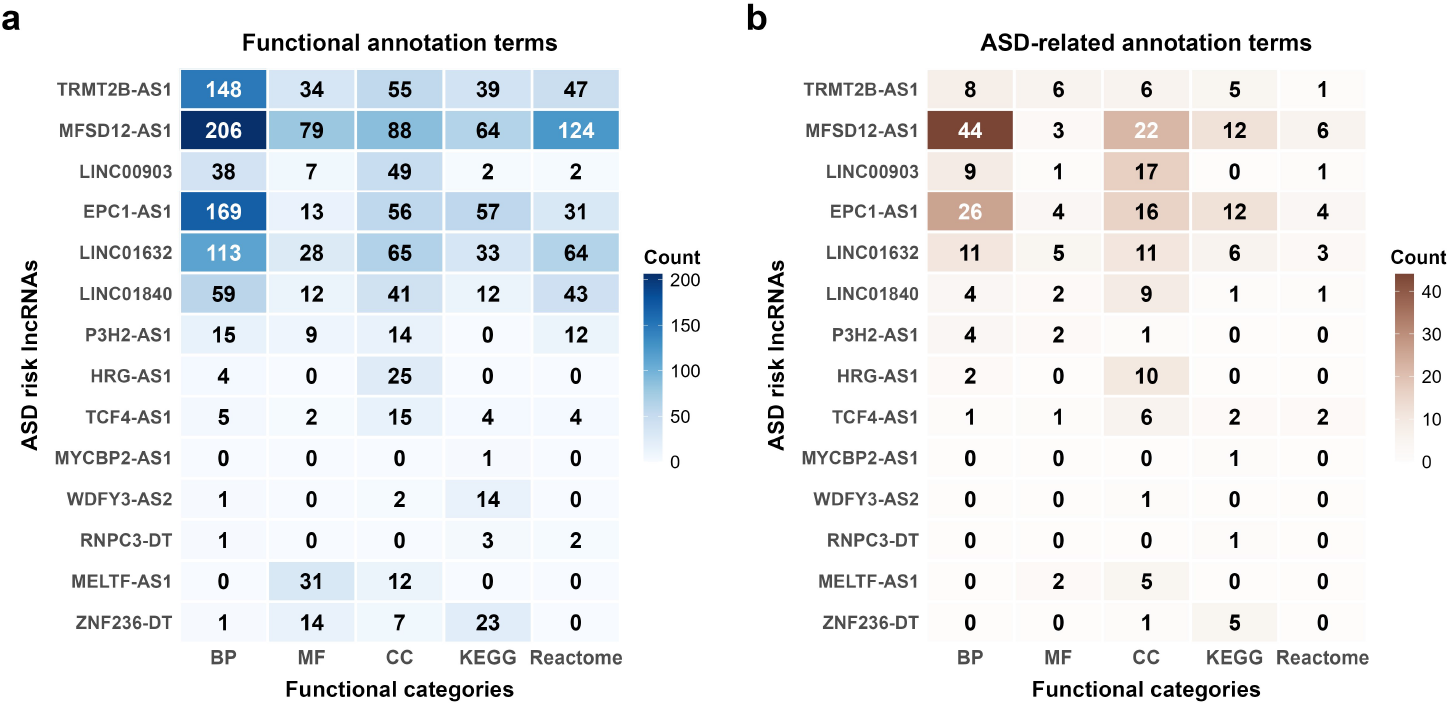
Functional enrichment analysis of 14 ASD risk lncRNAs. (a) Heat map of functional annotation terms of 14 ASD risk lncRNAs, and (b) Heat map of ASD-related annotation terms of 14 ASD risk lncRNAs.

### Potential lncRNA biomarkers enable robust classification of sex and cell types

Sex and cell type are two fundamental axes of biological heterogeneity in ASD (Lai et al. 2014; Iakoucheva et al. 2019b). Sex differences are crucial for understanding ASD prevalence, while cell-type-specific molecular disruptions are central to ASD’s neurodevelopmental pathology. Therefore, investigating potential lncRNA biomarkers across sex and cell type can reveal context-specific pathobiology and inform precise diagnostic strategies.

We identify distinct profiles of lncRNA biomarkers in ASD versus Normal contexts (**Fig. 5a, Supplemental Data 5**). For sex, a greater number of context-specific biomarkers (81) is found in ASD than context-enhanced ones (48), and a similar pattern is observed in Normal (128 vs. 56). In contrast, for cell type, context-enhanced biomarkers vastly outnumber context-specific ones in both ASD (268 vs. 27) and Normal (328 vs. 22). This divergence suggests that the biological underpinnings of ASD differ fundamentally across these two axes: sex effects may reflect more stable, developmental programming differences, whereas cell-type disruptions may indicate active, disease-progress-related alterations in specific cellular environments.

**Figure 5.**
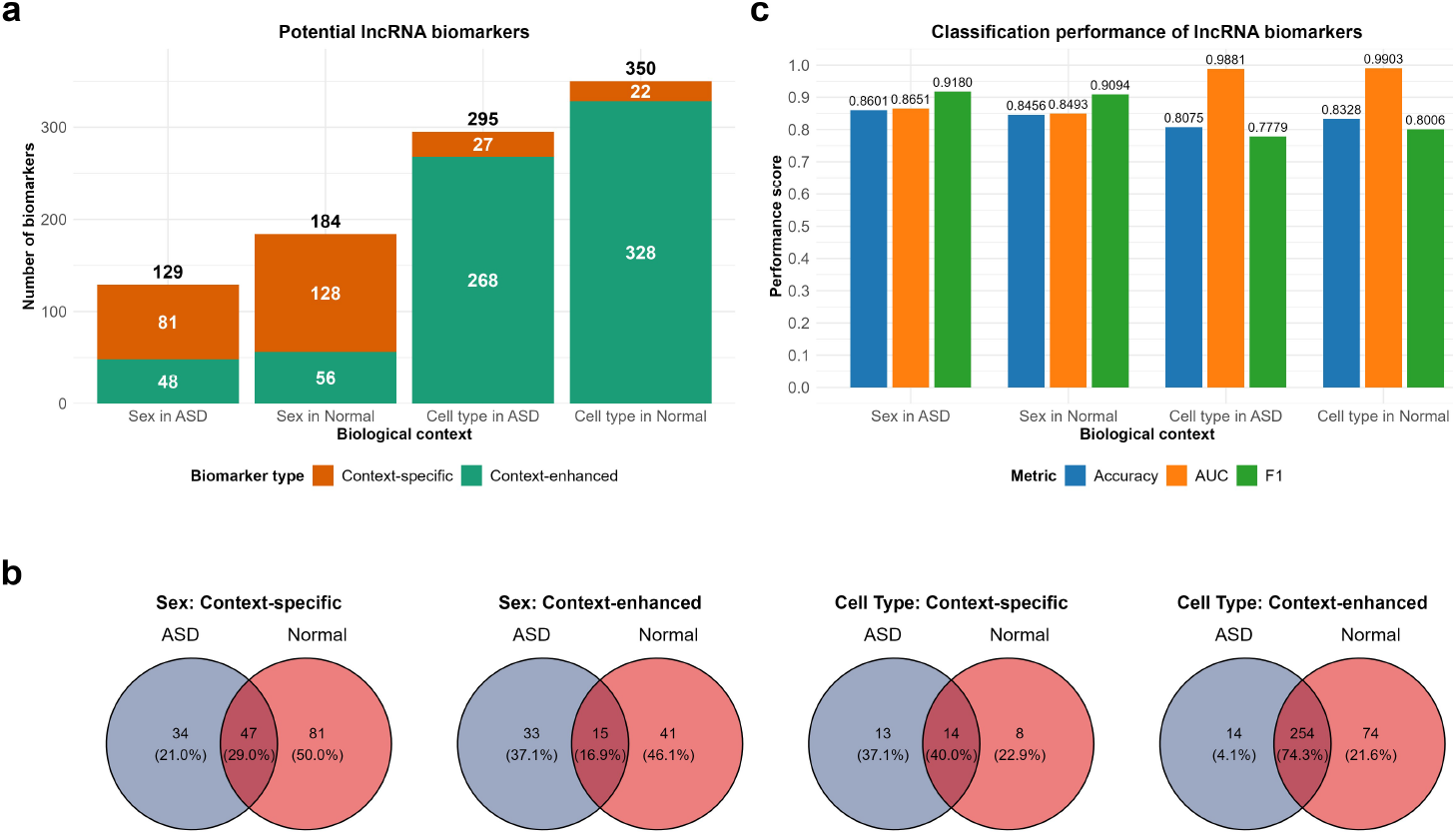
Potential lncRNA biomarkers. (a) Number of lncRNA biomarkers. (b) Venn diagrams of lncRNA biomarkers of sex and cell type. (c) Classification performance of lncRNA biomarkers in distinguishing sex and cell types.

A comparative analysis of shared lncRNA biomarkers between ASD and Normal reveals distinct patterns of overlap across sex and cell type (**Fig. 5b**). For sex, only a limited subset of context-specific (29.0%) and context-enhanced (16.9%) biomarkers are common between ASD and Normal. The small common set suggests that ASD substantially alters or employs different sex-biased lncRNA regulatory networks compared to Normal, pointing to a more pronounced rewiring of sex-specific molecular pathways in the disorder. In contrast, for cell type, a substantially larger proportion of context-specific (40.0%) and context-enhanced (74.3%) are shared between ASD and Normal. This result implies that cellular lncRNA regulatory architecture remains largely intact in ASD, with dysregulation often involving the enhancement of pre-existing cell-type-specific programs.

The classification performance of lncRNA biomarkers is evaluated using Accuracy, AUC, and F1 (**Fig. 5c**). In distinguishing sex within ASD, the lncRNA biomarker set achieves an Accuracy of 0.8601, AUC of 0.8651, and F1 score of 0.9180. For cell type classification in ASD, performance reach an Accuracy of 0.8075, AUC of 0.9881, and F1 score of 0.7779. Comparative analysis across contexts confirmed consistently high discriminatory power, with Accuracy, AUC and F1 score remaining above 0.80 for Normal. This result indicates that lncRNA biomarkers provide a reliable molecular tool for accurate sex and cell type discrimination in ASD and Normal contexts, supporting their potential for diagnostic applications.

### Immune-related lncRNAs are implicated in the immune regulation

Growing evidence implicates immune dysregulation in the pathophysiology of ASD, linking neuroinflammation and altered immune signaling to behavioral and synaptic anomalies (Garbett et al. 2008; Mead and Ashwood 2015). As an important layer of gene regulation, lncRNAs have been increasingly implicated in immune responses and neurodevelopmental processes (Atianand and Fitzgerald 2014; Parikshak et al. 2016). To investigate their potential role in ASD-related immune dysregulation, we identify two functionally distinct classes of immune-associated lncRNAs, including 29 context-specific and 11 context-enhanced lncRNAs. We further perform functional enrichment analysis to systematically characterize their involvement in immune regulation (**Supplemental Data 6**).

Functional enrichment analysis shows that 26 out of 29 context-specific lncRNAs are significantly associated with at least one BP, MF, CC, KEGG or Reactome functional term (**Fig. 6a**), and 15 lncRNAs (*AQP4-AS1, ATP6V0D1-DT, CBR3-AS1, DISC1FP1, FAM182B, HCG17, KCNQ1OT1, LINC00486, LINC01090, LINC02197, LY6E-DT, MAPT-AS1, MIR137HG, PRDM6-AS1, SNHG16*) are enriched in at least one functional term directly linked to immune regulation (**Fig. 6b**). Likewise, all 11 context-enhanced lncRNAs are enriched in at least one functional term (**Fig. 6c**), and four lncRNAs (*MAPKAPK5-AS1, MIR4300HG, UGDH-AS1, MAST3-AS1*) are involved in at least one functional term related to immune regulation (**Fig. 6d**). Additionally, previous studies support roles for *KCNQ1OT1* and *NEAT1* in immune modulation (Li et al. 2024; Tastan et al. 2025). Collectively, our findings suggest that the discovered 20 immune-related lncRNAs (16 context-specific and 4 context-enhanced immune lncRNAs) are implicated in the immune regulation of ASD, and may offer promising targets for further immune therapy.

**Figure 6.**
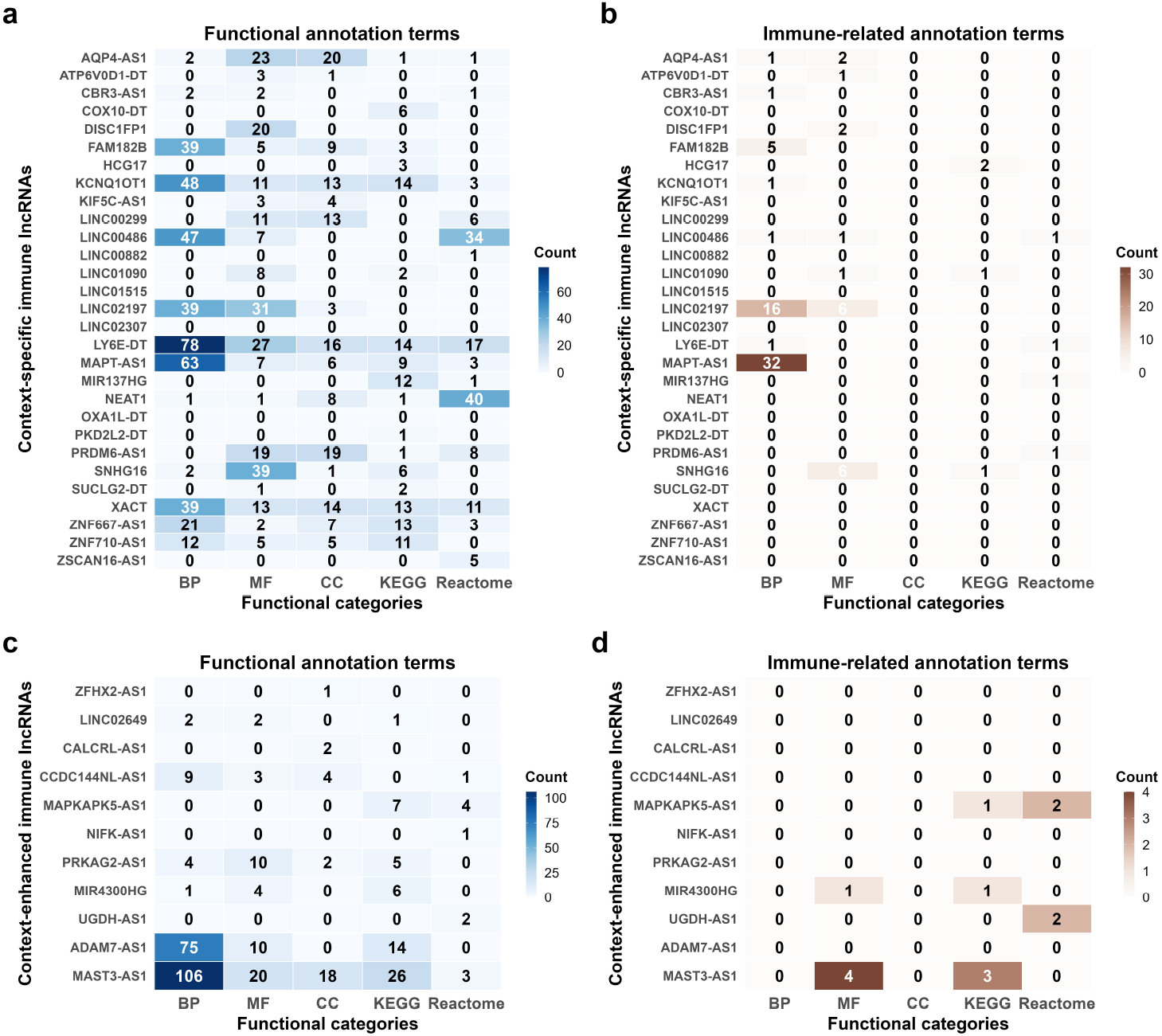
Functional enrichment analysis of 29 context-specific and 11 context-enhanced immune lncRNAs. (a) Heat map of functional annotation terms of 29 context-specific immune lncRNAs, (b) Heat map of immune-related annotation terms of 29 context-specific immune lncRNAs, (c) Heat map of functional annotation terms of 11 context-enhanced immune lncRNAs, and (d) Heat map of immune-related annotation terms of 11 context-enhanced immune lncRNAs.

### Cell-cell interaction networks diverge substantially between ASD and Normal

Dysregulation of cell-cell interaction networks represents a central pathological feature in ASD, reflecting altered neuroimmune crosstalk, glial support, and neuronal circuit coordination (Velmeshev et al. 2019). The comparative analysis of cell-cell interaction networks between ASD and Normal cortical tissues reveals substantial reorganization of intercellular signaling (Astorkia et al. 2022). To map these alterations, we integrate predicted lncRNA causal networks with putative ligand– receptor interactions to systematically construct cell-cell interaction networks for both ASD and Normal contexts (**Supplemental Data 7**).

In the ASD network, interactions involving Microglia show pronounced strength with multiple neuronal cell types (e.g., L23, INPV, L4) and other glial cells (e.g., ASTFB, ASTPP, OPC, Oligodendrocytes), suggesting an activated neuroimmune interface (**Fig. 7a**). Similarly, OPC and Oligodendrocytes exhibit enhanced connectivity, indicating dysregulated myelination or glial-neuronal support. In contrast, the Normal network displays more balanced and spatially structured interactions, with stronger engagement of ASTFB, NeuNRGNI and NeuNRGNII cell subtypes, which may reflect homeostatic astrocyte-neuron coupling (**Fig. 7b**).

**Figure 7.**
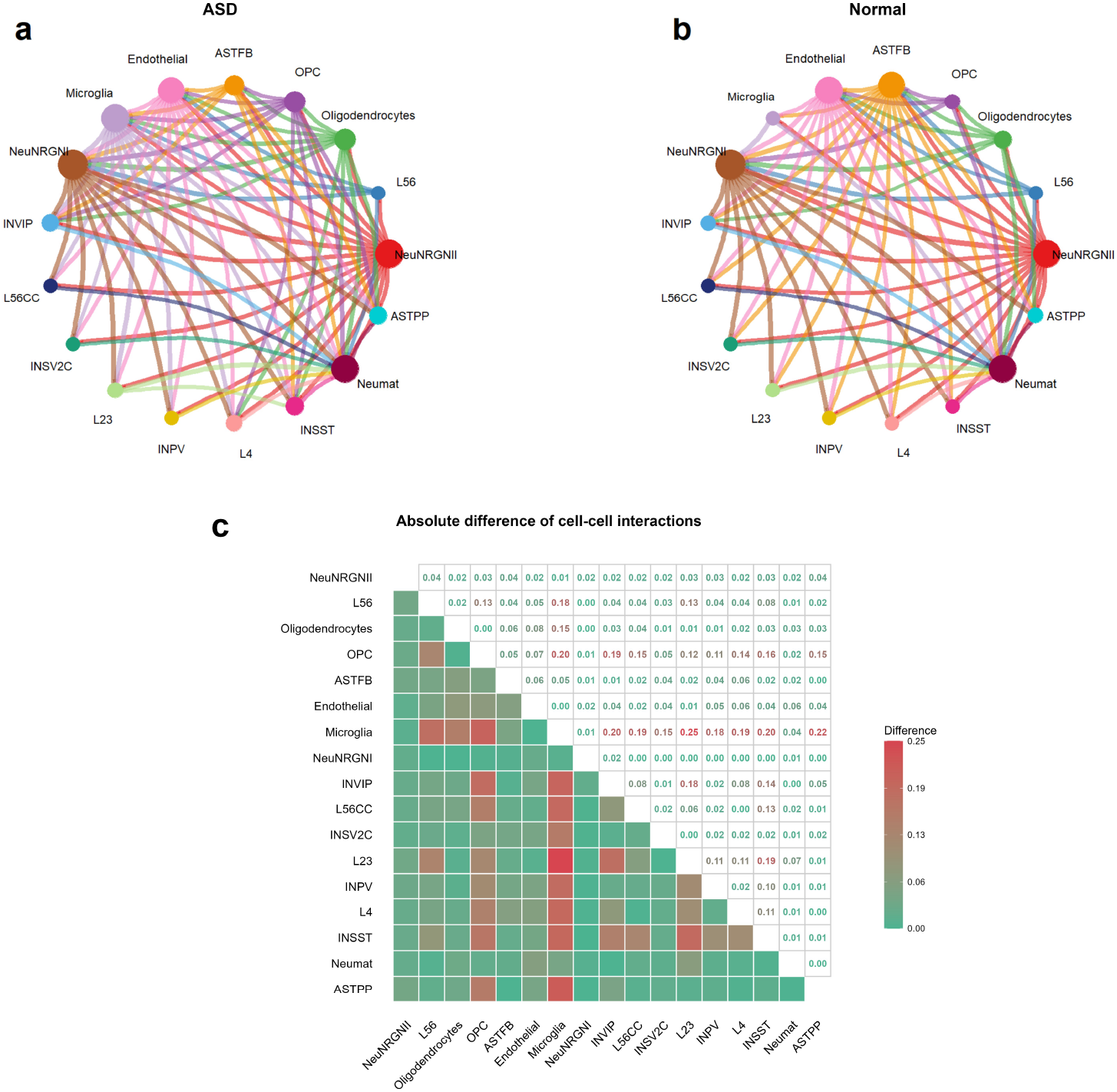
Cell-cell interactions in ASD and Normal. (a) Visualization of cell-cell interactions with a strength above a threshold of 0.60 in ASD. (b) Visualization of cell-cell interactions with a strength above a threshold of 0.60 in Normal. (c) Absolute difference of cell-cell interactions between ASD and Normal.

As shown in **Fig. 7c**, 20.59% of all cell-cell interactions (28 of 136) are substantially altered between ASD and Normal networks, with an absolute difference exceeding 0.10. Notably, Microglia show elevated differential engagement with several neuronal layers (e.g., L23, INPV, L4) and interneurons (e.g., INVIP, INSV2C), supporting the notion of immune-mediated synaptic modulation in ASD (Zhan et al. 2014). Furthermore, OPC interactions are broadly intensified, paralleling the evidence of white matter and oligodendrocyte dysfunction in ASD (Ohta et al. 2020; Travers et al. 2012). Subtle but consistent reductions are observed in ASTPP interactions with specific neuronal cell types, particularly inhibitory interneurons (e.g., INPV, INSST) and excitatory neurons in key cortical layers (e.g., L23, L4), suggesting a broad impairment in astrocytic modulation of synaptic efficacy and network integration. These findings imply that ASD pathophysiology arises not from isolated cellular deficits, but from a system-wide rewiring of the intercellular network. The divergence in interaction strength, particularly involving Microglia and OPC, underscores their potential role as cellular hubs in ASD pathophysiology.

## Discussion

In this work, we present Clean, a hidden causality-based framework designed to infer dynamic lncRNA causal regulation in ASD. Our analyses show that 14 ASD risk lncRNAs are significantly involved in ASD pathogenesis. Furthermore, lncRNA expression signatures enable robust classification of sex and cell type, highlighting their potential as predictive biomarkers. Our analyses identified 20 immune-related lncRNAs implicating neuroimmune dysregulation, alongside substantially altered cell-cell interaction networks in ASD. Despite these promising results, several limitations inherent to Clean need to be considered.

In SuperCell (Bilous et al. 2022), the construction of metacells involves three critical parameters (the number of nearest neighbors, the number of the top genes, and the graining level) that influence the balance between biological resolution and computational robustness. To prioritize fine-grained local structure, the number of nearest neighbors to build KNN (K-Nearest Neighbors) network is set to 5. The use of the top 2000 variable genes for dimensionality reduction is consistent with common practices in snRNA-seq data and supports efficient computation without substantial loss of biologically relevant information. A graining level of 30 is to define the approximate number of cells aggregated per metacell, and it achieves a practical trade-off between mitigating technical noise and preserving biological relevance. To ensure reliable downstream analysis (trajectory inference and hidden causal inference), future efforts will focus on enhancing the rigor of metacell partitioning in snRNA-seq data analysis.

For the application of trajectory inference to metacells, Clean apply a linear trajectory inference method SCORPIUS (Cannoodt et al. 2016), which assumes simplified, unidirectional transitions between cell states. While SCORPIUS is interpretable and efficient, it fails to capture complex biological processes such as branching events, cyclic dynamics, or convergent differentiation pathways in ASD. To more accurately delineate dynamic lncRNA causal regulation in ASD, Clean will use nonlinear trajectory inference approaches (e.g., Monocle (Trapnell et al. 2014), Slingshot (Street et al. 2018) and PAGA (Wolf et al. 2019)) to order metacells in future.

The parameter settings of hidden causal inference in Clean are carefully determined to balance computational feasibility and biological interpretability. The embedding dimension *E*=3 and time lag *τ* = 1 are selected in accordance with Takens’ theorem for state-space reconstruction (Sauer 2006). We select Euclidean distance metric due to its widespread applicability, and robustness in capturing distance within reconstructed state spaces. The number of steps ahead *h*=1 aims to infer immediate causal relationships, which is particularly meaningful given rapid responses in gene regulatory networks. A critical empirical parameter is the causal strength cutoff set at 0.60, used to identify significant lncRNA-mRNA causal relationships. This threshold seeks to minimize false positives while retaining biologically plausible causality. Future studies could benefit from applying adaptive cutoff strategies.

While data-driven hidden causal inference method Clean can identify temporal causality, it is often susceptible to false positives due to high-dimensionality and noise in snRNA-seq data. To enhance the accuracy and biological interpretability of inferred lncRNA causal regulatory networks, we incorporate the priori information of lncRNA targets predicted by LncTar. The combination of temporal causality and priori information can help distinguish direct regulation from indirect co-regulation. Future refinements could involve incorporating high-confidence lncRNA targets as priori information.

By providing single-nucleus resolution, snRNA-seq captures the full spectrum of cellular diversity in the brain through detailed transcriptional profiles. However, it may suffer from lower transcript capture efficiency and higher technical noise, particularly for lowly expressed lncRNAs. In contrast, bulk RNA sequencing offers high detection sensitivity for transcripts, especially low-abundance lncRNAs. In future, integrating snRNA-seq and bulk RNA sequencing data provides a powerful strategy to enhance the inference and functional interpretation of lncRNA causal regulation in autism.

The comprehensive evaluation of heterogeneity between networks is essential for quantifying structural and functional divergence in systems biology. In terms of lncRNA causal regulatory networks, we use Simpson index (Simpson 1949) to evaluate the heterogeneity between different brain biological contexts. Actually, the Jaccard index (Hancock 2014) and Lin’s method (Lin 1998) can also be used to characterize network heterogeneity. Applying a multi-metric index (combining Simpson index, Jaccard index and Lin’s method) would enhance interpretability and robustness in characterizing network heterogeneity, especially in ASD.

While it demonstrates exceptional power for classifying sex and cell type, the classification performance is moderate for diagnosis, brain region, and age. For example, the lncRNA biomarker set for distinguishing ASD from Normal achieves an accuracy of 0.5735, an AUC of 0.6065, and an F1 score of 0.5477. For classifying brain region, the classification model has an accuracy of 0.6589 (AUC: 0.6524, F1: 0.7755) within ASD and 0.6538 (AUC: 0.7147, F1: 0.7433) within Normal. Age prediction yields accuracies of 0.7020 (AUC: 0.6080, F1: 0.8210) in ASD and 0.6945 (AUC: 0.6765, F1: 0.8021) in Normal. This indicates that although potential lncRNA biomarkers can clearly distinguish basic biological categories (e.g., sex and cell type), their ability to differentiate systems-level traits (e.g., diagnosis, brain region, and age) is still limited.

Clean holds significant potential for extension to other regulatory ncRNAs, including microRNAs (miRNAs), circular RNAs (circRNAs) and Piwi-interacting RNAs (piRNAs). By refining priori information of ncRNA targets and tuning parameter settings to match ncRNA-specific causal regulatory dynamics, Clean could evolve into a unified causal inference platform for diverse regulatory ncRNAs. This would not only enhance our understanding of multi-layer ncRNA causal regulatory networks in ASD, but also facilitate the discovery of novel ncRNA biomarkers and therapeutic targets.

Taken altogether, we develop a computational framework Clean to delineate dynamic lncRNA causal regulation in ASD. Our findings reveal the context-specific heterogeneity of lncRNA causal interactions across diverse brain biological contexts, providing new insights into their regulatory roles. We anticipate that Clean will be widely applicable for uncovering lncRNA regulation mechanisms in complex neurodevelopmental disorders.

Taken altogether, we develop a computational framework Clean to delineate dynamic lncRNA causal regulation in ASD. Our findings reveal the context-specific heterogeneity of lncRNA causal interactions across diverse brain biological contexts, providing new insights into their regulatory roles. We anticipate that Clean will be widely applicable for uncovering lncRNA regulation mechanisms in complex neurodevelopmental disorders.

## Methods

### Single-nucleus RNA sequencing data in autism

We obtained the largest single-nucleus RNA sequencing (snRNA-seq) data of cortical tissue including the anterior cingulated (ACC) and prefrontal cortex (PFC) regions from patients with autism, accessed in Sequence Read Archive (SRA) with accession number PRJNA434002 (Velmeshev et al. 2019). We utilized gene annotation information from HGNC (HUGO Gene Nomenclature Committee) (Seal et al. 2023) to further divide RNA transcripts into lncRNAs and mRNAs. Given that low-expression genes might lack biological significance, we restricted our analysis to genes demonstrating expression levels above the mean expression value across all cells. In total, we have identified 813 lncRNAs and 5,133 mRNAs exhibiting high expression levels across 104,559 cells (including 52,003 autism cells and 52,556 normal cells). Based on the known expression of cell type markers (Velmeshev et al. 2019), the 52,003 autism cells and 52,556 normal cells are annotated into 17 cell types, including fibrous astrocytes (AST-FB), protoplasmic astrocytes (AST-PP), oligodendrocyte precursor cells (OPC), layer 2/3 excitatory neurons (L2/3), layer 4 excitatory neurons (L4), layer 5/6 corticofugal projection neurons (L5/6), layer 5/6 cortico-cortical projection neurons (L5/6-CC), parvalbumin interneurons (IN-PV), somatostatin interneurons (IN-SST), SV2C interneurons (IN-SV2C), VIP interneurons (IN-VIP), maturing neurons (Neu-mat), NRGN-expressing neurons (Neu-NRGN-I), NRGN-expressing neurons (Neu-NRGN-II), endothelial cells, microglia cells and oligodendrocytes.

### Metacells building

By merging gene expression of similar autism and normal brain cells, Clean uses SuperCell (Bilous et al. 2022) to build metacells of snRNA-seq data. For SuperCell, three parameters (the number of nearest neighbours, the number of the top genes, and the graining level) need to set. Empirically, the number of nearest neighbors to build KNN (K-Nearest Neighbors) network is set to 5, and the number of the top variable genes to use for dimensionality reduction is set to 2000. Since the graining level (i.e., level of size reduction between the snRNA-seq and the metacell data) is generally recommended between 10 and 50, Clean selects five graining levels (10, 20, 30, 40 and 50) and then use four metrics (including purity, compactness, separation, and inner normalized variance) to perform quality controls on metacells by using MetacellAnalysisToolkit (Bilous et al. 2024). Here, purity is the proportion of the most abundant cell type within the metacell, compactness is the variance of the components within the metacell, separation is the distance to the closest metacell, and inner normalized variance is the mean-normalized variance of gene expression within the metacell. Higher values of purity and separation, and lower values of compactness and inner normalized variance, indicate better metacell partitioning performance. In terms of separation and inner normalized variance, larger graining levels have better metacell partitioning. However, in terms of purity and compactness, lower graining levels perform better in metacell partitioning. For a compromise, Clean sets the graining level to 30 in this work.

### Trajectory inference

Given the metacell data of each data slice, Clean applies SCORPIUS (Cannoodt et al. 2016) to perform linear trajectory inference on metacells. SCORPIUS reduces the dimensionality of the metacell data, and returns coordinates of the metacells represented in an 2-dimensional space. Based on the identified 2-dimensional space, SCORPIUS further orders brain metacells of each data slice along a linear trajectory. Consequently, we have obtained 48 pseudo time-series metacell data for hidden causal inference.

### Priori information of lncRNA targets

To enhance the accuracy of dynamic lncRNA causal regulation, we apply a sequence-based method LncTar (Li et al. 2015) to identify putative lncRNA targets. We obtain the sequences of lncRNAs and mRNAs from the Ensembl (Dyer et al. 2025) database (version 115) via the biomaRt (Durinck et al. 2009) R package, prioritizing the longest transcript isoform per gene. LncTar predicts lncRNA–mRNA interactions by calculating the minimum free energy of the joint secondary structure formed through base pairing between two RNA transcripts. For large-scale identification of lncRNA targets, we utilize a parallelized implementation of LncTar. In this study, we applied a normalized binding free energy (ndG) threshold of ≤ -0.1 to infer significant lncRNA-mRNA interactions. In total, we have obtained a list of 105,649 lncRNA-mRNA interactions as priori information of dynamic lncRNA causal regulation (**Supplemental Data 8**).

### Dynamic lncRNA causal regulation

Causal networks are abundant in biological processes, and the hidden nature of causality is positive, negative, or dark in complex biological systems (Stavroglou et al. 2019, 2020). To quantify positive, negative, and dark causality in the pseudo time-series metacell data, the PC (Pattern Causality) algorithm (Stavroglou et al. 2019, 2020) is introduced for hidden causal inference.

Let *R* and *G* represent the time-series data for a lncRNA and an mRNA, respectively. For *R* and *G*, PC creates two shadow attractors, *M*_*R*_ and *M*_*G*_, respectively, by finding the optimal pair (*E,τ*). Here, *E* denotes the embedding dimension of the shadow attractor, and *τ* is the time lag to construct a shadow attractor.

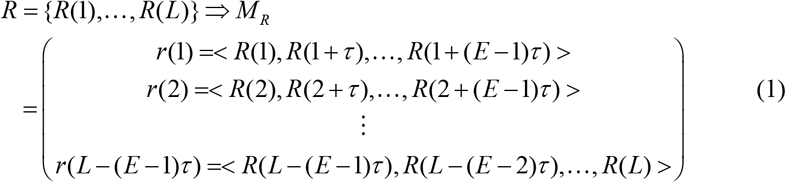

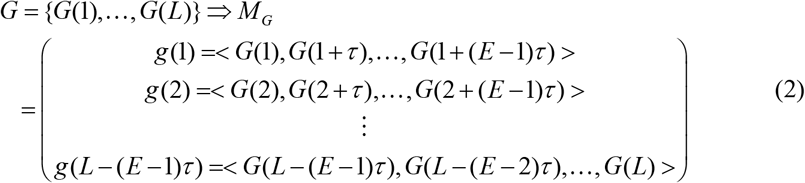

Where *L* is the length of the time-series data, *r* (*t*) and *g* (*t*) are the vectors of *M*_*R*_ and *M*_*G*_ respectively corresponding to the state at time *t*. Among all vectors in *M*_*R*_ and *M*_*G*_, the distance matrices *D*_*R*_ and *D*_*G*_ are calculated:

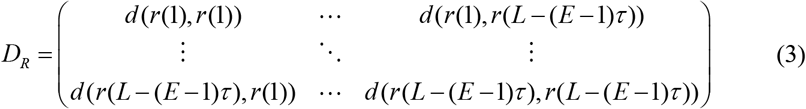

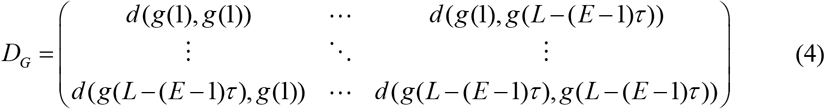

For each point *r*(*t*) in *M*_*R*_, its *E*+1 nearest neighbours *NN*_*r* (*t*)_ is found. From the *E* +1 nearest neighbours, the projected time indices 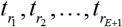 by projecting *t*_1_,*t*_2_, …,*t*_*E*+1_ ahead by *h* steps are:

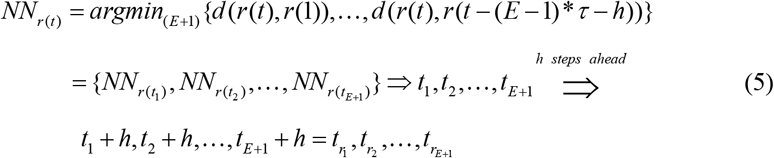

The distance of the projected *E* +1 nearest neighbours from *r*(*t*) and *g*(*t*):

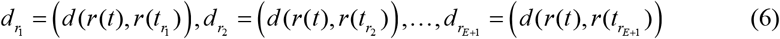

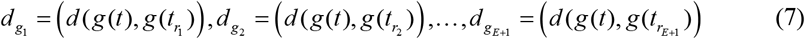

The future pattern *P*_*g* (*t* +*h*)_ of *g*(*t* + *h*) at time *t* + *h* is estimated:

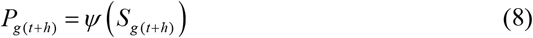

Where the mapping function*Ψ* (·) is a pattern extraction function that maps the space

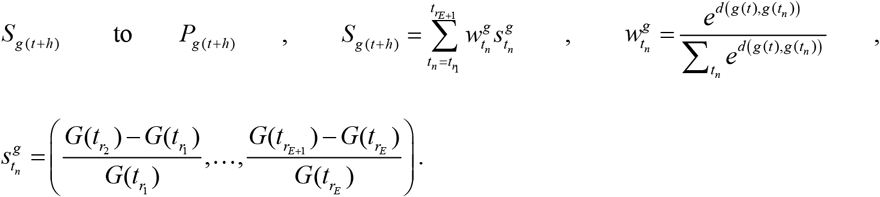

The current pattern *P*_*r* (*t*)_ of *r* (*t*) at time *t* is calculated:

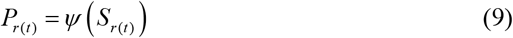

Where the mapping function*Ψ* (·) is a pattern extraction function that maps the space

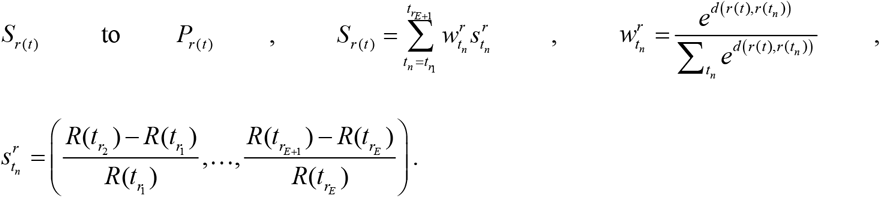

The strength of pattern causality from *R* to *G* at every time step *t* is:

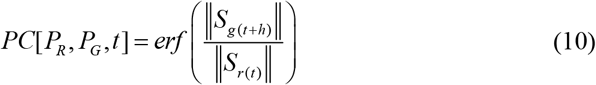

Where ‖*S*_*g* (*t* +*h*)_ ‖ and ‖*S*_*r* (*t*)_ ‖ represent the norms of *S*_*g* (*t* +*h*)_ and *S*_*r* (*t*)_ respectively, the error function 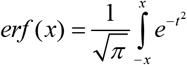.

At time *t*, positive, negative, and dark causality are represented as *P*(*t*), *N*(*t*), and *D*(*t*), respectively. Since multiple causality types cannot coexist, only one type of causal pattern exist at a time *t*. The causality type with the largest causal strength is regarded as the final causal pattern from *R* to *G*.

In this work, Clean utilizes patterncausality (Wang et al. 2024) R package to calculate positive, negative and dark causality between lncRNAs and mRNAs. When using the patterncausality tool, the embedding dimension *E* is set to 3, the time lag *τ* is 1, the distance metric used is Euclidean, and the number of steps ahead *h* is 1. To accelerate the speed of hidden causal inference, Clean uses parallel computing with 48 CPU cores to infer positive, negative and dark causality between lncRNAs and mRNAs. For identifying significant lncRNA-mRNA causal relationships, the cutoff of positive, negative or dark causal strength is empirically set to 0.60. The causality type with the largest causal strength is regarded as the final causal pattern from a lncRNA to an mRNA. Given 48 pseudo time-series metacell data spanning five brain biological contexts (i.e. diagnosis, region, age, sex and cell type), Clean constructs 48 lncRNA causal regulatory networks in total.

### Heterogeneity analysis

Given two lncRNA causal regulatory networks *Net*_*i*_ and *Net*_*j*_, the heterogeneity *Het*(*Net*_*i*_, *Net*_*j*_) between *Net*_*i*_ and *Net*_*j*_ is defined as:

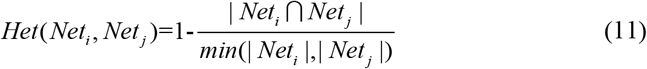

Where | *Net*_*i*_ ∩ *Net* _*j*_ | is the number of common lncRNA-mRNA causal relationships between *Net*_*i*_ and *Net*_*j*_, *min*(| *Net*_*i*_ |,| *Net*_*j*_ |) denotes the number of lncRNA-mRNA causal relationships in the smallest network between *Net*_*i*_ and *Net*_*j*_. *Het*(*Net*_*i*_, *Net*_*j*_) ranges from 0 to 1. A higher value of *Het*(*Net*_*i*_, *Net*_*j*_) implies that *Net*_*i*_ and *Net*_*j*_ have greater heterogeneity with each other. In terms of lncRNA causal regulatory networks, we use *Het*(*Net*_*i*_, *Net*_*j*_) to evaluate the heterogeneity between different brain biological contexts (i.e. diagnosis, region, age, sex and cell type).

### ASD risk lncRNAs identification

Differential expression analysis is commonly employed to identify ASD lncRNAs. However, for single-cell RNA sequencing data, differential expression analysis is challenged by excessive zeros, normalization, donor effects, and cumulative biases (Wu et al. 2025). Additionally, differential expression analysis focuses on lncRNA expression levels, and ignores their downstream targets. To address these limitations, we develop a network-based approach to identify ASD risk lncRNAs by analyzing differential causal networks between ASD and Normal. For each lncRNA, we quantify network rewiring by calculating the total number of rewired causal edges (i.e., sum of gained and lost edges). The risk score (RS) and significance *p*-value for each lncRNA *i* is computed as follows.

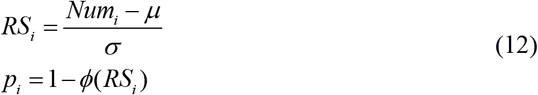

Where *Num*_*i*_ is the number of rewired causal edges for lncRNA *i, μ* represents the mean number of rewired causal edges across all lncRNAs, *σ* denotes the standard deviation, and *ϕ* is the cumulative distribution function (CDF) of the standard normal distribution.

We define lncRNAs with statistically significant risk scores (*p* < 0.05) as ASD risk lncRNAs, characterized by markedly disrupted causal regulatory patterns in ASD.

### Potential lncRNA biomarkers prediction

Hub lncRNAs are highly connected nodes in lncRNA causal regulatory networks, and are often regarded as promising biomarkers. Here, we identify hub lncRNAs by applying the cumulative Poisson distribution probability:

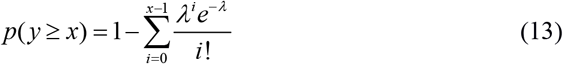

where 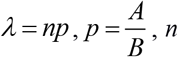 is the number of genes (including lncRNAs and mRNAs), *A* is the number of lncRNA-mRNA causal relationships in a lncRNA causal regulatory network, and *B* is the number of all possible lncRNA-mRNA causal pairs. Smaller *p*-value of a lncRNA indicates that the lncRNA is more likely to be a hub. Here, the cutoff of the *p*-value is set to 0.05.

In this work, cell type biomarkers include two types: cell-type-specific and cell-type-enhanced. Moreover, sex biomarkers also include two types: sex-specific and sex-enhanced. Hub lncRNAs identified exclusively in a single cell type or sex are cell-type-specific or sex-specific biomarkers. A hub lncRNA is a cell-type-enhanced or sex-enhanced biomarker if its degree in a specific cell type or sex is at least two-fold greater than its average degree across all other cell types or sex.

### Immune-related lncRNA detection

Among 17 major cell types in the brain, microglia are the resident immune cells of the central nervous system, and play a critical role in the pathogenesis of ASD (Salter and Stevens 2017; Tetreault et al. 2012). By modulating gene expression in immune cells such as microglia, lncRNAs play crucial regulatory roles in the neuroimmune system (Salta and De Strooper 2017). In this work, we define that hub lncRNAs specifically identified in microglia under either ASD or Normal conditions are context-specific immune lncRNAs. Additionally, a hub lncRNA identified in microglia is also considered as an immune lncRNA (i.e., a context-enhanced immune lncRNA) if its degree in ASD is at least two-fold higher or lower than its degree under normal condition. Similar to lncRNA biomarkers prediction, we also identify hub lncRNAs by applying the cumulative Poisson distribution probability.

### Functional enrichment analysis

We conduct functional enrichment analysis using clusterProfiler (Wu et al. 2021) to assess the involvement of ASD risk lncRNAs in the pathogenesis of ASD, and the implication of immune-related lncRNAs in the immune regulation. Functional enrichment analysis is conducted for each ASD risk lncRNA using its rewired targets between ASD and Normal, whereas for each immune-related lncRNA, both rewired and shared targets between ASD and Normal are used. In this work, the functional enrichment analysis cover Gene Ontology (GO) (Ashburner et al. 2000) including Biological Process (BP), Molecular Function (MF), and Cellular Component (CC), as well as the Kyoto Encyclopedia of Genes and Genomes (KEGG) (Kanehisa and Goto 2000) and Reactome (Milacic et al. 2024) pathways. ASD risk and immune-related lncRNAs are significantly enriched in GO, KEGG, and Reactome terms (adjusted *p* < 0.05, adjusted by Benjamini-Hochberg method).

### Classification analysis

To evaluate the diagnostic performance of the identified lncRNA biomarkers, we employ a classification analysis using Support Vector Machine (SVM) (Cortes and Vapnik 1995) implemented in the e1071 (Meyer et al. 2024) R package. Separate classification tasks are performed for binary (sex) and multiclass (cell type) predictions within both ASD and Normal cohorts. The discriminative capacity of each model is assessed using 10-fold cross-validation. Model performance for both binary and multiclass classification is evaluated using three standard metrics: Accuracy, Area Under the receiver operating characteristic Curve (AUC), and F1-score. Here, Accuracy is the proportion of correct predictions among the total number of cases, AUC measures the overall ability of a model to distinguish between classes across all classification thresholds, and F1-score is the harmonic mean of precision and recall. Higher values of Accuracy, AUC, and F1-score indicate better classification performance.

### Cell-cell interaction network construction

In this work, we integrate predicted lncRNA causal regulatory networks with putative ligand–receptor interaction networks to assess intercellular interaction strength between different cell types. The ligand–receptor interactions are compiled from five databases: CellPhoneDB (Troulé et al. 2025), CellChatDB (Jin et al. 2021), CellTalkDB (Shao et al. 2021), Cellinker (Zhang et al. 2021b), and CellCall (Zhang et al. 2021a). We retained only those ligand–receptor pairs involving mRNAs present in the predicted lncRNA causal networks for each cell type. In terms of the combined networks ( *CN*_*i*_ and *CN* _*j*_ for cell types *i* and *j* respectively), the interaction strength *IS* (*i, j*) between the two cell types is calculated as:

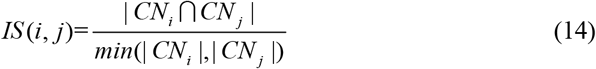

Where | *CN*_*i*_ ∩ *CN* _*j*_ | indicates the number of shared interactions between *CN*_*i*_ and *CN* _*j*_ *min*(| *CN*_*i*_ |,| *CN* _*j*_ |) represents the number of interactions in the minimum network between *CN*_*i*_ and *CN* _*j*_. The value of *IS* (*i, j*) ranges from 0 to 1, where 0 indicates no interaction and 1 indicates perfect interaction.

### Method execution

Each execution of Clean on single-nucleus RNA sequencing data is performed in a separate task. Each task is allocated 48 CPU cores of Intel(R) Xeon(R) Platinum 8375C CPU at 2.90 GHz, and one R session is opened for each task.

## Code availability

Clean is released under the GPL-3.0 License, and is available at https://github.com/zhangjunpeng411/Clean/.

## Data access

The largest single-nucleus RNA sequencing (snRNA-seq) data are accessed from Sequence Read Archive (SRA) with accession number PRJNA434002. All data supporting the findings of the current study are listed in Supplementary Data and our GitHub website (https://github.com/zhangjunpeng411/Clean/).

## Author contributions

J.Z. conceived the idea of this work. X.W. and C.Z. refined the idea. J.Z. designed and performed the experiments. Z.H. and S.F. participated in the design of the study and performed the statistical analysis. J.Z., X.W. and C.Z. drafted the manuscript. All authors revised the manuscript. All authors read and approved the final manuscript.

## Supplemental Data

**Supplemental Data 1**. Summary of 48 data slices of autism snRNA-seq data.

**Supplemental Data 2**. Number of positive, negative and dark causality.

**Supplemental Data 3**. Risk scores of lncRNAs.

**Supplemental Data 4**. Functional enrichment analysis of 14 ASD risk lncRNAs.

**Supplemental Data 5**. Potential lncRNA biomarkers of sex and cell types.

**Supplemental Data 6**. Enrichment analysis of 40 immune-related lncRNAs.

**Supplemental Data 7**. Cell-cell interaction strength in ASD and Normal.

**Supplemental Data 8**. Priori information of lncRNA targets predicted by LncTar.

